## Supporting Information for "Mono-mix strategy enables comparative proteomics of a cross-kingdom microbial symbiosis"

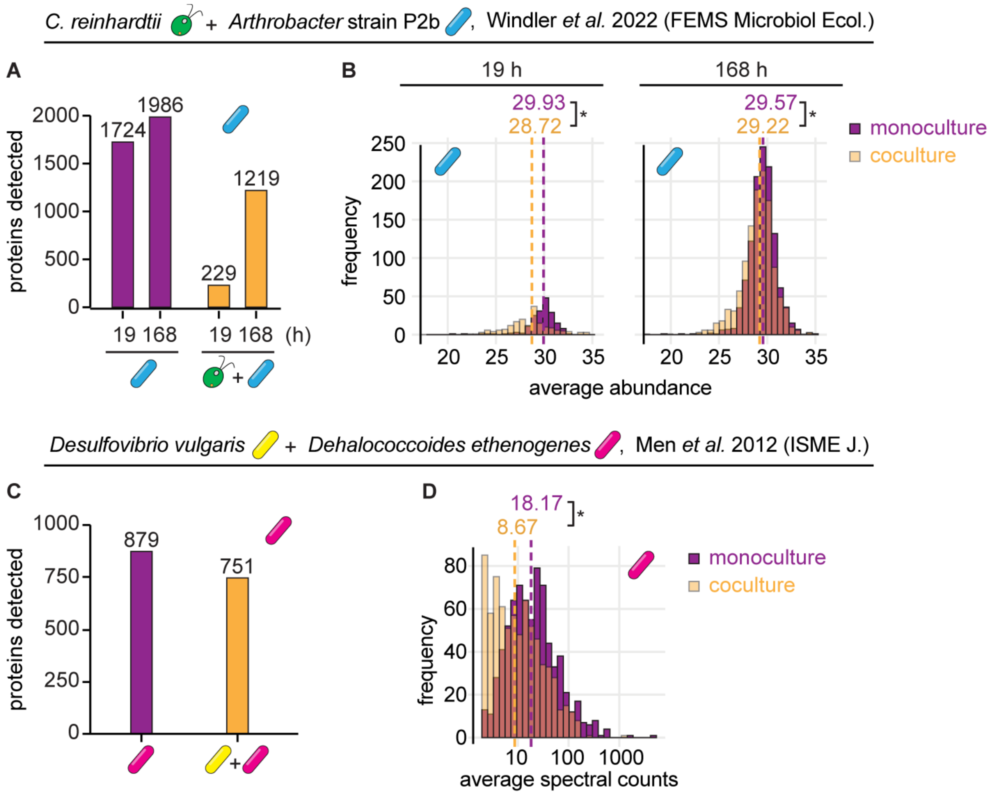


**S1 Fig. Bacterial protein detection is decreased in other coculture proteomic studies; related to Fig 1.**

- **(A)** Number of unique *Arthrobacter* proteins with at least two peptides detected in at least one biological replicate of monocultures (purple) and cocultures with *C. reinhardtii* (orange) 19 h and 168 h after inoculation by LC-MS/MS proteomics reported in Windler *et al.* 2022 [21].
- **(B)** Distribution of *Arthrobacter* protein abundances in monocultures (purple) and cocultures with *C. reinhardtii* (transparent orange) for proteins detected in both conditions reported in Windler *et al.* 2022 [21]. When distributions overlap, the color is muddied. Median values are shown above the dashed lines. Asterisks next to the medians indicate significant differences between the indicated distributions by a Wilcoxon rank-sum test ($p < 0.05$).
- **(C)** Number of unique *Dehalococcoides ethenogenes* proteins detected in at least two biological replicates of monocultures (purple) and cocultures with *Desulfovibrio vulgaris* (orange) by LC-MS/MS proteomics reported in Men *et al.* 2012 [48].
- **(D)** Distribution of *D. ethenogenes* protein abundances in monocultures (purple) and cocultures with *D. vulgaris* (transparent orange) for proteins detected in both conditions reported in Men *et al.* 2012 [48], presented as in (**B**).

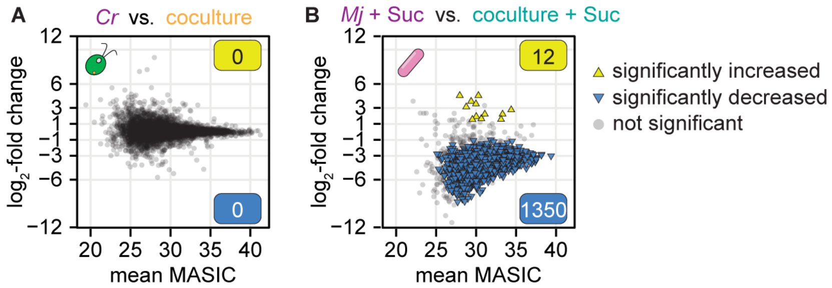


**S2 Fig. Global decrease in bacterial protein detection complicates differential expression analysis, especially prior to data normalization; related to Fig 1.** Triplicate continuous-light-grown monocultures and cocultures with and without sucrose were collected for LC-MS/MS proteomics.

- **(A)** Changes in unnormalized algal protein abundances in coculture relative to monoculture. Significant differences (triangles) were defined as those where $|\log_2$-fold change$| > 1$, *p-adj.* $< 0.05$ from a Student's t-test of the unnormalized MASIC values, and the mean MASIC value was greater than the limit of quantitation in both the coculture and the monoculture.
- **(B)** Changes in unnormalized bacterial protein abundances presented as in (**A**).

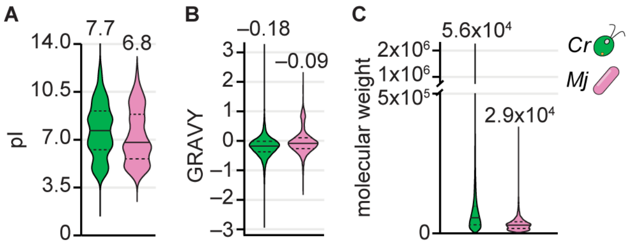


**S3 Fig. Physicochemical characteristics of proteins encoded by *C. reinhardtii* and *M. japonicum* vary considerably; related to Fig 2.**

The physicochemical characteristics of *C. reinhardtii* and *M. japonicum* proteins were assessed *in silico* according to Shi *et al.* 2024 [36].

- **(A)** Distribution of isoelectric points (pI) of proteins encoded by *C. reinhardtii* (green) and *M. japonicum* (pink). Median values are represented by the solid horizontal lines and listed above the violins, and quartiles are represented by dashed lines.
- **(B)** Distribution of hydropathy (grand average of hydropathy, GRAVY) of proteins encoded by *C. reinhardtii* (green) and *M. japonicum* (pink), presented as in (**A**).
- **(C)** Distribution of molecular weights of proteins encoded by *C. reinhardtii* (green) and *M. japonicum* (pink), presented as in (**A**).

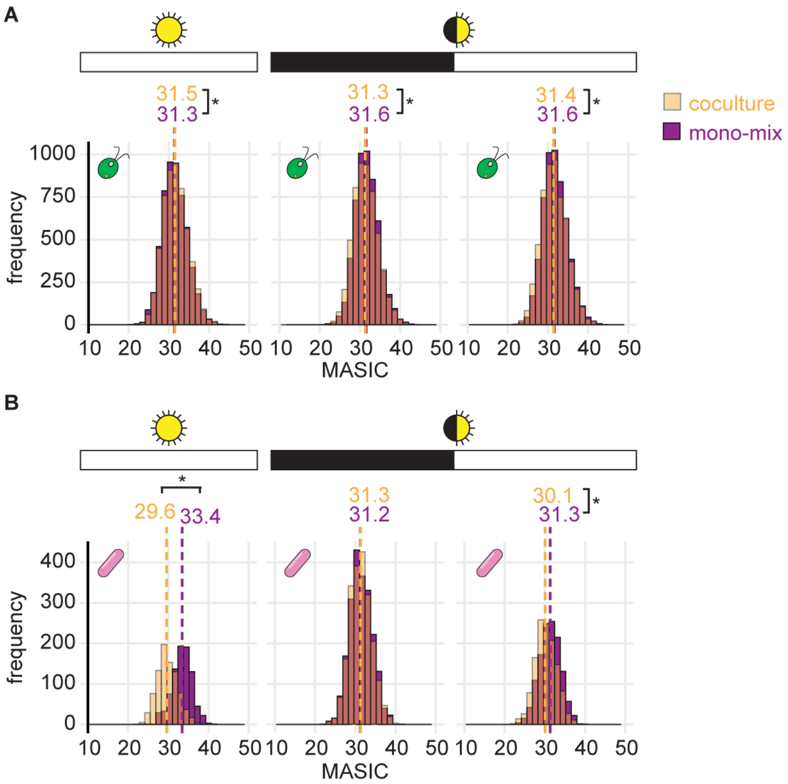


**S4 Fig. Distribution of unnormalized protein abundances in the mono-mix experiment; related to Fig 2.**

Triplicate cocultures of *C. reinhardtii* and *M. japonicum* were grown in parallel with triplicate monocultures of *C. reinhardtii* and *M. japonicum* with 150 µg/ml sucrose. Then, the *M. japonicum* monocultures were added to the *C. reinhardtii* monocultures to achieve a "mono-mix" control with a similar bacteria-to-algae ratio as the coculture. Cultures were grown in either continuous or diurnal light. Continuous light cultures were collected 36 h after inoculation, and the diurnal light cultures were collected at the end of the night (36 h after inoculation) and the end of the day (48 h after inoculation).

- **(A)** Distribution of unnormalized abundances of algal proteins in cocultures (transparent orange) and mono-mix controls (purple) grown in continuous light (sun icon) or diurnal light (eclipsed sun icon) collected at the end of the dark or light phases (black or white bars, respectively). When distributions overlap, the color is muddied. The MASIC values were averaged across the three biological replicates. Median values are shown above the dashed lines. Asterisks next to the medians indicate significant differences between the indicated distributions by a Wilcoxon rank-sum test ($p < 0.05$).
- **(B)** Distribution of unnormalized abundances of bacterial proteins presented as in (**A**).

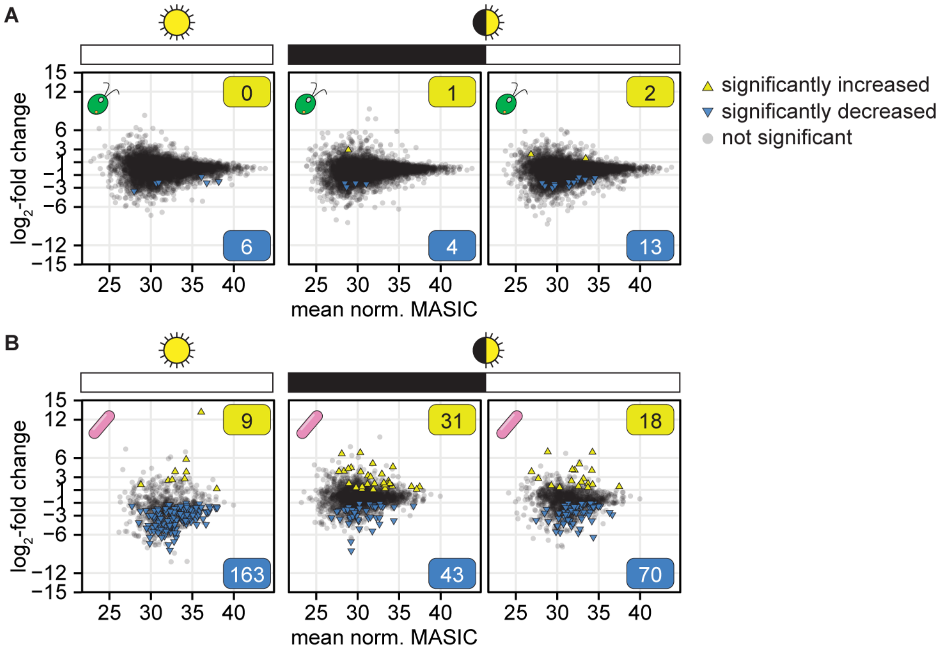


**S5 Fig. Differential expression analysis using the mono-mix strategy may be sensitive to differences in organism relative abundance, even after normalization; related to Figs 2 and 3.**

**(A)** Changes in quantile-normalized algal protein abundances in coculture relative to mono-mix controls when grown in continuous light (sun icon) or diurnal light (eclipsed sun icon) and collected at the end of the dark or light phases (black or white bars, respectively). Significant differences (triangles) were defined as those where |$\log_2$-fold change| > 1, *p-adj.* < 0.05 from a Student's t-test of the quantile-normalized MASIC values, and the mean MASIC value was greater than the limit of quantitation in both the coculture and the monoculture.

**(B)** Changes in quantile-normalized bacterial protein abundances in coculture relative to mono-mix controls presented as in (**A**).

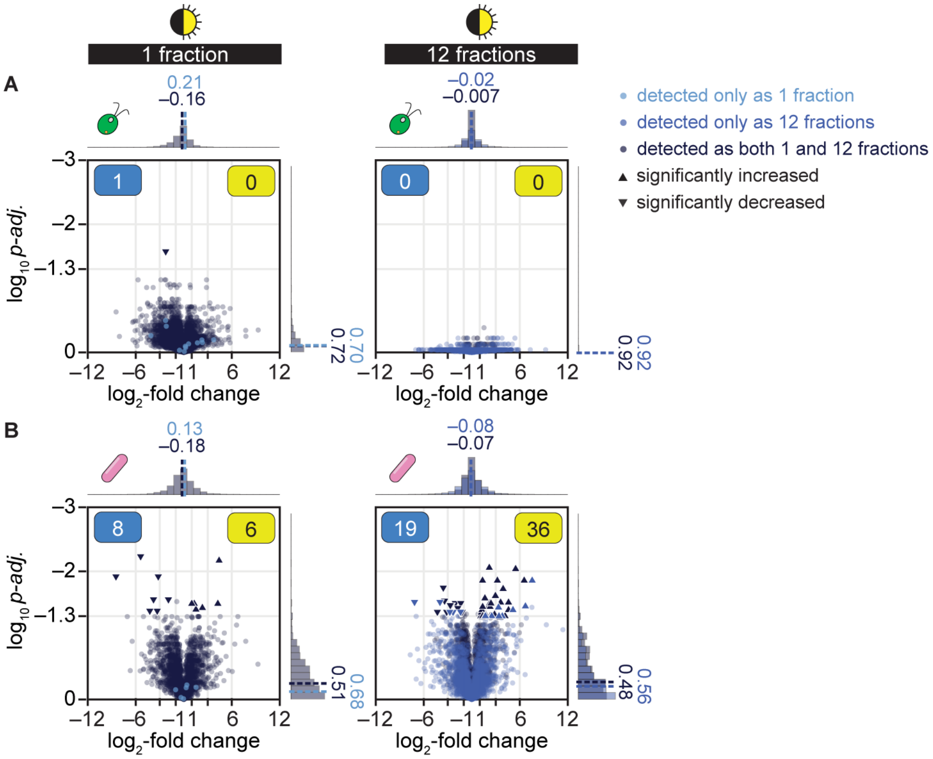


**S6 Fig. Sample fractionation increases the signal-to-noise for proteomic differential expression analysis of proteins from coculture relative to mono-mix controls; related to Fig 5.**

**(A)** Changes in quantile-normalized *C. reinhardtii* protein abundance in coculture relative to mono-mix controls at the end of the night when samples were analyzed as 1 (left) or 12 fractions (right). Points are colored based on their detection criteria: lighter blue indicates that the protein only met the criteria when the sample was analyzed as either 1 or 12 fractions, navy blue indicates that the protein met the criteria when the sample was analyzed both as 1 and as 12 fractions. Significant differences (triangles) were defined as those where $|\log_2\text{-fold change}| > 1$, *p-adj.* $<$ 0.05 from a Student's t-test of the quantile-normalized MASIC values, and the mean MASIC value was greater than the limit of quantitation in both the coculture and the mono-mix. Yellow and blue boxes indicate the number of significantly increased and decreased proteins, respectively. Histograms show the distribution of $\log_2$-fold changes (*x* axes) and of $\log_{10}$ *p-adj.* (*y* axes) for proteins that only met the detection criteria when the sample was analyzed as either 1 or 12 fractions (lighter blue) and for proteins that met the detection criteria when the sample was analyzed both as 1 and as 12 fractions (navy blue); median values are shown above the dashed lines.

**(B)** Changes in quantile-normalized *M. japonicum* protein abundance in coculture relative to mono-mix controls at the end of the night when samples were analyzed as 1 (left) or 12 fractions (right) presented as in (**A**).

**S1 Table: Sample metadata for the proteomics samples; related to S2, S3, and S5 Tables.**

**S2 Table: Algal and bacterial protein abundances under various interaction regimes; related to Fig 1 and S2 Fig.**

**S3 Table: Algal and bacterial protein abundances under various interaction and light regimes determined using the mono-mix strategy; related to Figs 2–3 and S4–S5 Figs.**

**S4 Table: Significant differences in protein abundance in coculture relative to mono-mix controls under various light regimes; related to Fig 3 and S5 Fig.**

**S5 Table: Algal and bacterial protein abundances under various interaction regimes at the end of the night determined using the mono-mix strategy with and without sample fractionation; related to Figs 4 and 5.**

**S6 Table: Significant differences in protein abundance in coculture relative to mono-mix controls at the end of the night, as determined with or without sample fractionation; related to Fig 5 and S6 Fig.**
